# Droplet-based microfluidics platform for investigation of protoplast development of three exemplary plant species

**DOI:** 10.1101/2025.08.19.671073

**Authors:** Paulina Marczakiewicz-Perera, Traud Winkelmann, Michael Köhler, Jialan Cao

**Affiliations:** Institute for Chemistry and Biotechnology, Dept. of Physical Chemistry and Microreaction Technology, Technische Universität Ilmenau, 98693 Ilmenau, Germany; Institute of Plant Genetics, Section Reproduction and Development, Leibniz Universität Hannover, 30419 Hannover, Germany

## Abstract

Microfluidic technologies offer powerful tools for miniaturized and highly controlled biological experiments, yet their application in plant research remains underexploited. In this study, we present a droplet-based microfluidic platform tailored for the encapsulation and cultivation of plant protoplasts, enabling long-term observation of cell development at nearly single-cell resolution. Protoplasts isolated from leaves of *Nicotiana tabacum, Brassica juncea*, and *Kalanchoe daigremontiana* were used to evaluate the platform’s suitability across diverse plant species. Our results demonstrate species-dependent responses to microfluidic cultivation, with tobacco protoplasts showing the highest viability. The system permits dynamic tracking of cell fate within individual droplets and supports the quantification of stochastic and concentration-dependent responses to chemical stimuli. Using tobacco protoplasts, we further investigated the effect of low concentrations of cytokinins (BAP) and auxins (NAA) for the early protoplast culture, up to the first division. Low concentrations (20–80 µg·L^−1^) significantly enhanced cell survival and cell growth, while higher doses did not yield additional benefits. This work underscores the potential of droplet-based microfluidics as a high-resolution, low-volume platform for protoplast-based assays and dose-response screening, with applications across diverse plant biotechnology studies.

## 1. Introduction

Droplet-based microfluidics has emerged as a powerful tool for miniaturized, high-throughput biological experimentation, enabling precise control over cellular environments, single-cell resolution, and massively parallel assays. By compartmentalizing reactions or individual cells into nanoliter-sized droplets, these platforms facilitate highly reproducible, scalable studies that would be difficult or impossible using conventional bulk methods. While this technology has seen wide adoption in biomedical and microbial research [1,2], its application in plant biology, particularly for fragile, wall-less cells such as protoplasts, remains limited and does not include prolonged cultivation [3–5].

Protoplasts, isolated plant cells without their cell walls, are inherently sensitive and highly responsive to microenvironmental cues. These properties make them both technically challenging and biologically informative as a model system for studying cell viability, division, and differentiation. Beyond their utility in basic research, protoplasts serve as a versatile platform in plant biotechnology, enabling somatic hybridization, transient expression assays, and stable transformation through regeneration into whole plants. They are widely used in DNA delivery and genome editing workflows, providing a direct route to study gene function and engineer new traits without the constraints imposed by cell walls [6–8]. Traditionally, protoplast research has relied on static culture systems that offer limited control over spatial and temporal conditions, making it difficult to study heterogeneous responses at the single-cell level or in dynamic conditions [9–11].

Here, we present an improved droplet-based microfluidic platform, previously demonstrated for successful application in microspore embryogenesis investigation [12]. This system allows for the encapsulation, tracking, and longitudinal observation of individual protoplasts or small cell clusters within discrete aqueous droplets with precise control over droplet volume and chemical composition. Our primary objectives were to (i) assess the suitability of this microfluidic approach for maintaining plant protoplast viability over extended culture periods, (ii) evaluate its capacity to monitor early cell division events at nearly single-cell resolution, and (iii) explore the influence of low growth regulator concentrations on developmental outcomes.

By using protoplasts from three distinct plant species: *Nicotiana tabacum, Brassica juncea* and *Kalanchoe daigremontiana* as a test case, we demonstrate the platform’s capacity to reveal species-specific sensitivities, growth dynamics, and viability trends, as well as seek to as well as seek to identify technical constraints and optimization strategies for long-term culture. This work not only advances technical capabilities for plant single-cell analysis but also establishes a framework for broader applications in regenerative plant biotechnology and precision phenotyping.

## 2. Materials and methods

### 2.1. Plant culture

Plant material used in this study included *Kalanchoe daigremontiana* plantlets, *Nicotiana tabacum* cultivar Samsun NIC 422, and *Brassica juncea* CR 2664 seeds obtained from the gene bank of the IPK Gatersleben, Germany. Seeds were surface disinfected by immersion in 70% ethanol for 30 sec, followed by treatment with 2% sodium hypochlorite (NaOCl) containing a drop of Tween-20 for 5 min. They were then rinsed three times with sterile distilled water for 5 min each. *Kalanchoe* plantlets were disinfected using the same NaOCl solution for 10 min, omitting the ethanol step. Following disinfection, seeds and plantlets were transferred to Murashige and Skoog (MS) medium containing 30 g·L^−1^ sucrose and solidified with 4 g·L^−1^ gelrite. Germination and cultivation were conducted in a controlled environment at 24–25 °C with a 16 h light / 8 h dark photoperiod and a light intensity of approximately 5.0×10^−5^ mol·m^−2^·s^−1^. Tobacco plants were subcultured every 3–4 weeks. Mustard plants were freshly sown for each experiment and used at 2–3 weeks of age. *Kalanchoe* plants were propagated vegetatively every 3 months.

### 2.2. Isolation of protoplasts

For standard droplet cultivation experiments, protoplasts were isolated via enzymatic digestion from the top leaves of tobacco, mustard and *Kalanchoe* plants of various age (tobacco: 3-4 weeks old, mustard: 2 weeks old, *Kalanchoe*: 3 months old) following procedure described by [13]. Briefly, leaves were cut into small pieces on sterile sheets of paper and incubated in preplasmolysis solution for 1 h. The solution was then replaced with 5 mL of enzyme mixture (1.6% cellulase and 0.8% macerozyme in BNE9 solution, previously stirred, centrifuged, and filter sterilised), and samples were incubated for 15–17 hours at 27 °C in darkness, followed by an additional 30 min of gentle shaking at room temperature (36 rpm). The digested material was filtered through 100 µm sieves (Greiner Bio-One GmbH) into 13 mL round-bottom centrifuge tubes. An equal volume of 20% sucrose solution (5 mL) was added, mixed gently by inversion, and overlaid with 1.5 mL of washing solution to form two phases. After centrifugation at 760 g for 5 min, protoplasts collected at the interphase were transferred to fresh tubes, washed twice with 5 mL of washing solution (centrifuged at 700 g for 5 min each), and finally resuspended in 2 mL of 8pm7 cultivation medium. Based on cell counts using an Anvajo cell counter, the initial protoplast culture was adjusted to a density of 1.5–4 × 10^5^ cells·mL^−1^.

For growth regulators dose response experiments protoplasts were isolated from young tobacco leaves using a modified version of the protocol described by [14]. The initial disinfection step differed: leaves were halved and incubated in 2% PPM (Plant Preservative Mixture, Gentaur GmbH) for 1–2 hours at room temperature with gentle shaking, then rinsed with sterile water. All subsequent steps—including cutting, enzymatic digestion (using 0.25% cellulase and 0.25% macerozyme in F-PIN solution), filtration, two-phase separation, and washing – were performed as described above, with minor adjustments: digestion was carried out for 14 hours at 26 °C in darkness, followed by 30 minutes of shaking at room temperature. Isolated protoplasts were resuspended in 4 mL of F-PCN medium without growth regulators at a concentration of 1.6 × 10^5^cells·mL^−1^. For establishing a concentration range of hormonal treatments, F-PCN medium containing 4 mg·L^−1^ BAP and NAA was prepared and serially diluted as needed. The composition of all media used during protoplast isolation and culture is listed in *Supplementary Table S1*.

### 2.3. Microfluidic setup for cultivation in microdroplets

The experimental setup was based on previously described microfluidic systems [12,15], with several modifications (Fig. 1). Droplets were generated using a 6-port manifold, with flow rates precisely controlled by a multi-syringe pump (NEMESYS, Cetoni GmbH, Germany). For the standard droplet generation system, 1 mL glass syringes (SETonic GmbH, Germany) were used to deliver the cultivation medium and any effector solutions, while a 2.5 mL glass syringe was used for the carrier phase (PP9). All elements of the system were connected through fluorinated ethylene propylene (FEP) tubing with an inner diameter (ID) of 0.5 mm and outer diameter (OD) of 1.6 mm. This configuration ensured precise flow control.

**Figure 1.**
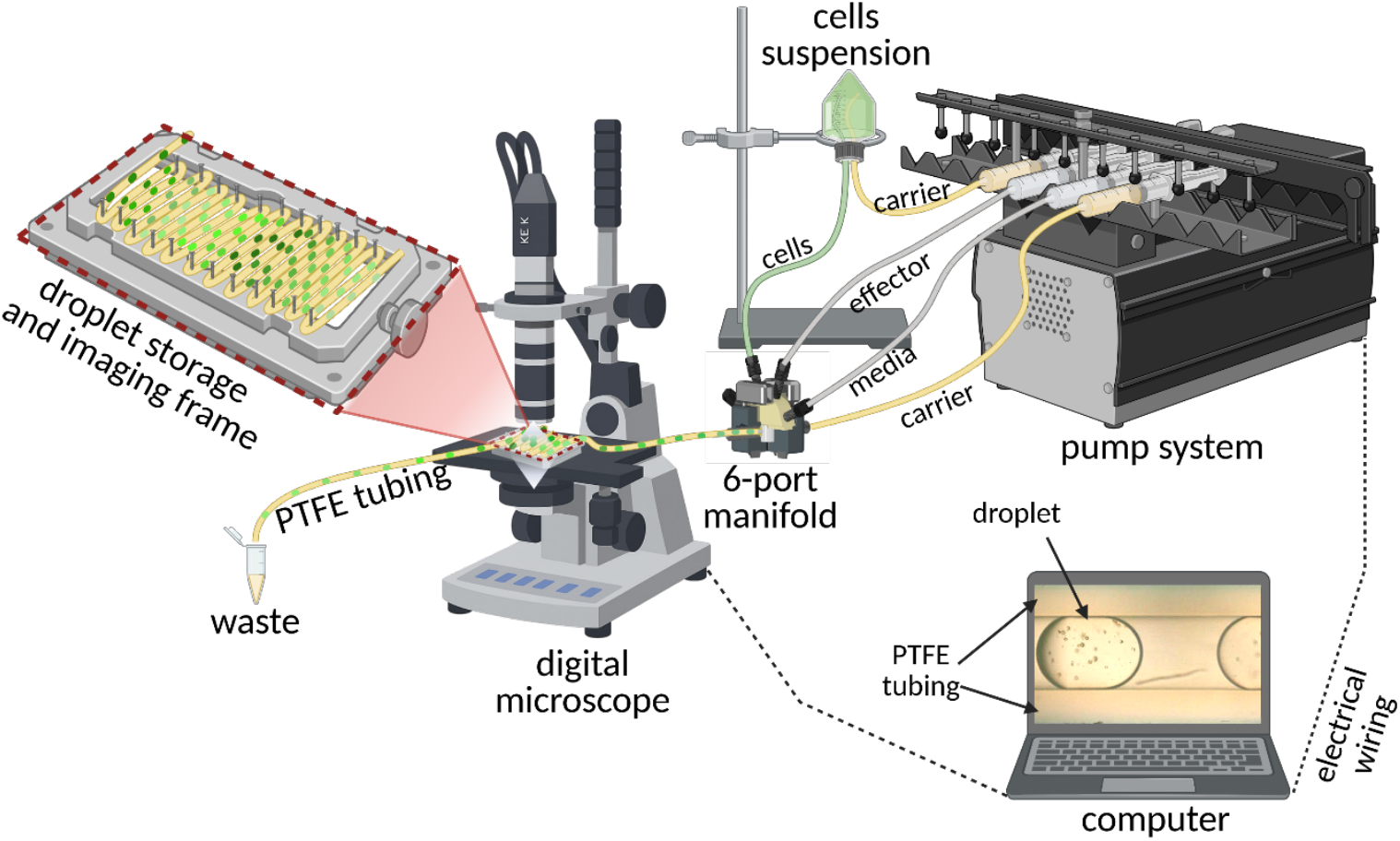
Experimental microfluidic setup. Droplets were generated using a 6-port manifold, enabling simultaneous delivery of up to two effectors. Protoplasts were introduced indirectly via a cell suspension container, where the carrier phase (PP9) pushed them toward the generator, reducing mechanical stress. Flow of all solutions was precisely controlled by a multi-syringe pump system. Droplet formation was monitored via a light camera. Culture occurred in sealed incubation tubing placed on a frame for imaging purposes using a digital microscope. Illustration created with Biorender.com.

To minimize mechanical stress on the protoplasts, cell suspensions were not directly injected. Instead, PP9 was introduced into a glass container holding the protoplast suspension, displacing the contents via a smaller polytetrafluoroethylene (PTFE) tube (ID 0.3 mm) toward the droplet generator. The remaining two syringes were used to deliver cell culture medium (for adjusting cell density) and, when applicable, specific effectors. Standard flow rates were approximately 20 µL·min^−1^ for the aqueous phase and 30 µL·min^−1^ for the continuous PP9 phase. Following droplet formation (typically 120–300 nL per droplet), they were directed into PTFE incubation tubing with thinner walls (ID 0.5 mm, OD 1.0 mm) to improve air perfusion. After formation, droplets were incubated in the sealed tubes in darkness at 24°C.

### 2.4. Reference cultivation in Petri dishes

In parallel with droplet-based experiments, bulk protoplast cultures were prepared as controls under identical conditions. Freshly isolated protoplasts were transferred into 6 cm Petri dishes containing 4 mL of respective cultivation medium (8pm7 or F-PCN) or into 12-well plates with 1 mL of medium per well, supplemented with varying concentrations of the tested effectors. Cell density was adjusted and maintained in the range of 1–2 × 10^4^ cells·mL^−1^.

### 2.5. Growth characterization and data processing

Protoplast cultures were monitored at regular intervals (every 1–3 days) using a digital microscope (Keyence VHX-5000) to assess cell morphology and detect the onset of cell division. In the dose response screening experiment, respective droplets could be found back based on their initial position in the incubation tubing. For quantitative analysis, the number and diameter of cells within individual droplets were measured manually using ImageJ (Fiji) software.

### 2.6. Statistics and reproducibility

Each experiment was conducted in duplicate, with 5-15 droplets analysed per condition in each run. Statistical analyses were performed in Jamovi (version 2.7.5).

For *Kalanchoe* protoplast size measurements, differences across cultivation days (0, 2, 5, 7) were assessed using the Kruskal–Wallis test, as normality was violated (Shapiro–Wilk, p < 0.001). Post-hoc for the growth regulator dose–response experiment with *Nicotiana tabacum* protoplasts, data were first tested for normality using the Shapiro–Wilk test. With only a few isolated deviations, most groups did not significantly depart from a normal distribution. Therefore, repeated measures ANOVA was applied to assess the effects of Time (days 1, 3, 5, 7; within-subject factor), Concentration (0, 20, 50, 80, 150 µg·L^−1^ NAA + BAP; between-subject factor), and their interaction on protoplast growth dynamics. Normality of residuals was evaluated with the Shapiro–Wilk test, and the Greenhouse–Geisser correction was applied where sphericity was violated. Changes in protoplast number and diameter (expressed relative to day 0) were analyzed accordingly. Estimated marginal means (EMMs) with 95% confidence intervals were calculated to visualize significant effects. Statistical significance was defined as α = 0.05.

## 3. Results and discussion

### 3.1. Comparison of protoplast development in bulk culture and in droplet culture

To evaluate whether the process of loading and cultivation in microfluidic droplets itself influences protoplast growth and survival, a standard bulk culture on Petri dishes was carried out in parallel with each experimental replicate under identical conditions. As shown in Figure 2, protoplast types exhibited typical developmental patterns in bulk culture, with initial cell divisions observable at around day 5 to 6. Tobacco protoplasts showed a tendency to form aggregates (Fig. 2c, bottom right). For both tobacco and *Kalanchoe* protoplasts, development proceeded similarly in droplets and in bulk culture, with droplet-cultured tobacco protoplasts displaying slightly larger diameters and a more rounded morphology compared to those in Petri dishes. In contrast, while mustard protoplasts divided normally in bulk, nearly all cells cultured in droplets were non-viable by day 6 (Fig. 2b), indicating a marked sensitivity to the mechanical stress associated with droplet formation. As this variability among plant species in protoplast behavior was observed during droplet cultivation, it is essential to first address the key limitations of this approach before discussing the detailed progression of cell growth and development. One of the primary concerns in microfluidic systems is the generation of shear stress, which varies depending on channel geometry and flow rates, and can lead to mechanical damage and reduced cell viability [16,17]. Sensitivity to shear stress differs not only between tissue types – such as leaves and hypocotyls – but also between species, influenced by factors such as genotype and cell wall characteristics [18].

**Figure 2.**
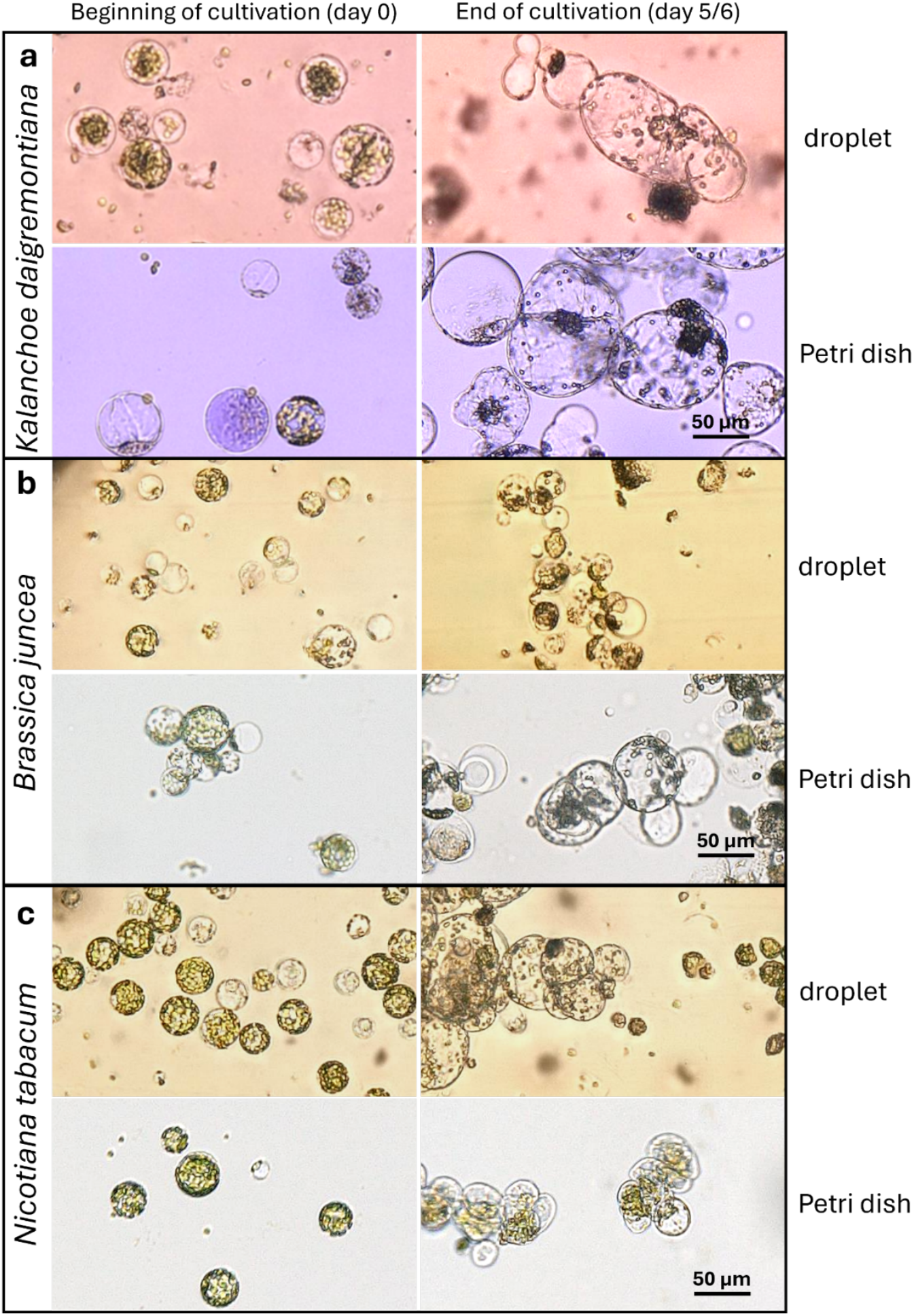
Comparison of protoplast appearance and development in droplets (top images in each panel) and in Petri dish bulk culture (bottom images), from culture initiation to the expected timing of first cell divisions (around day 5–6): (**a**) *Kalanchoe*; mustard; (**c**) tobacco.

### 3.2. Droplet culture generation and cell fate throughout cultivation

As mentioned in the previous section, the droplet generation process had a notable impact on the development of certain protoplasts. Here, we discuss in detail the behavior and developmental fate of protoplasts derived from the three studied plant species.

Mustard protoplasts exhibited pronounced stress responses immediately following droplet generation. Many cells displayed cytoplasmic condensation with organelle aggregation, accompanied by a significant number of cellular debris in the surrounding medium. While a few protoplasts remained viable during the initial days of cultivation, by day 5 the majority had become visibly shrunken and non-viable (Fig. 3), indicating poor tolerance to the shear stress during droplet formation.

**Figure 3.**
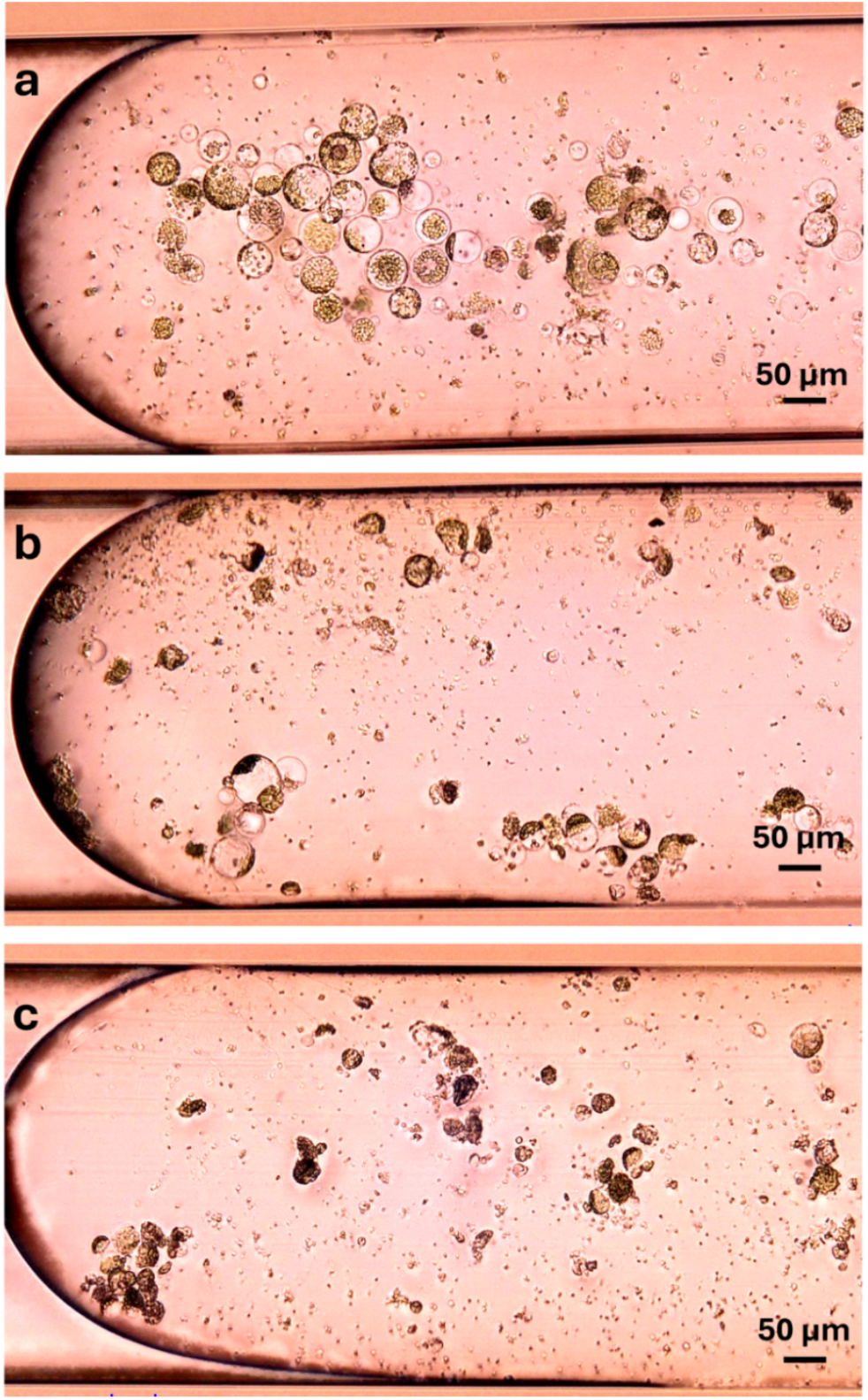
*Brassica juncea* leaf protoplasts at different time points during droplet cultivation: (a) day 0 – intact protoplasts were successfully introduced into droplets, although accompanied by cellular debris; (b) day 1 – some protoplasts maintained integrity, while many exhibited shrinkage and collapse; (c) day 5 – nearly all protoplasts had died. Images represent different droplets at each time point.

Tobacco protoplasts initially showed variability during isolation, but the final suspensions used for droplet loading were relatively uniform in morphology and free of large debris. These protoplasts tolerated the droplet generation process well, showing high integrity and minimal signs of mechanical damage or early cytotoxicity. Within the first two days of cultivation, the cells began to aggregate – particularly near interfaces – possibly as a protective response to the microenvironmental factors such as nutrient gradients or surface tension. This clustering complicated quantitative assessments at the later time points. Nevertheless, by day 6, numerous viable cells had undergone their first division, and microcallus formation was observed (Fig. 4d), suggesting successful adaptation and developmental progression within the droplet-based culture. Our results match the reports of durability of tobacco leaf protoplasts in microfluidic-based approaches [19,20].

**Figure 4.**
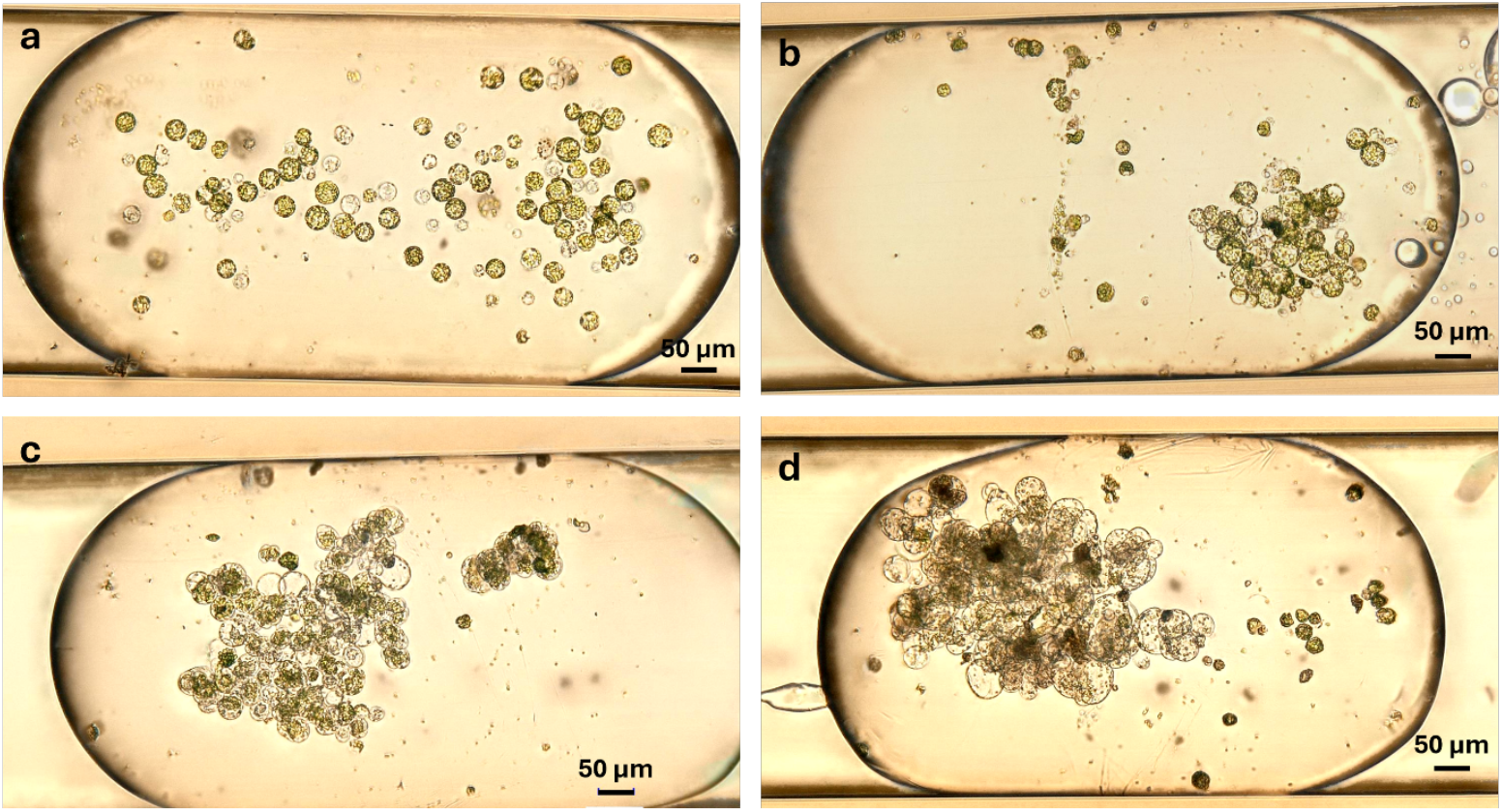
*Nicotiana tabacum* leaf protoplasts at different time points during droplet cultivation: (a) day 0 - healthy protoplasts successfully encapsulated; (b) day 1 – protoplasts remained viable and began coming into contact with each other, (c) day 3 – cells aggregated in clusters, (d) day 6 – first cell division took place, microcalluses were formed, it was impossible to count separate cells. Images represent different droplets at each time point.

*Kalanchoe* protoplasts exhibited pronounced heterogeneity in both size and internal composition immediately following isolation. Some cells exhibited intense violet pigmentation due to the accumulation of anthocyanins, while others contained only chloroplasts or were devoid of visible pigments. Post-isolation, these protoplasts had a tendency to aggregate and float in the standard cultivation medium, which hindered uniform dispersion in the loading reservoir and resulted in uneven distribution across droplet segments. Despite this, the majority of cells appeared intact immediately after droplet loading, with minimal debris present. By day 2, distinct cell enlargement was evident in a subset of protoplasts, while others remained static or displayed signs of cell death. By day 5, evidence of cell division was observed in some cells, whereas others continued to increase in size without undergoing division (Fig. 5). This divergent behavior suggests differences in developmental competence across the population, possibly influenced by the original tissue origin.

**Figure 5.**
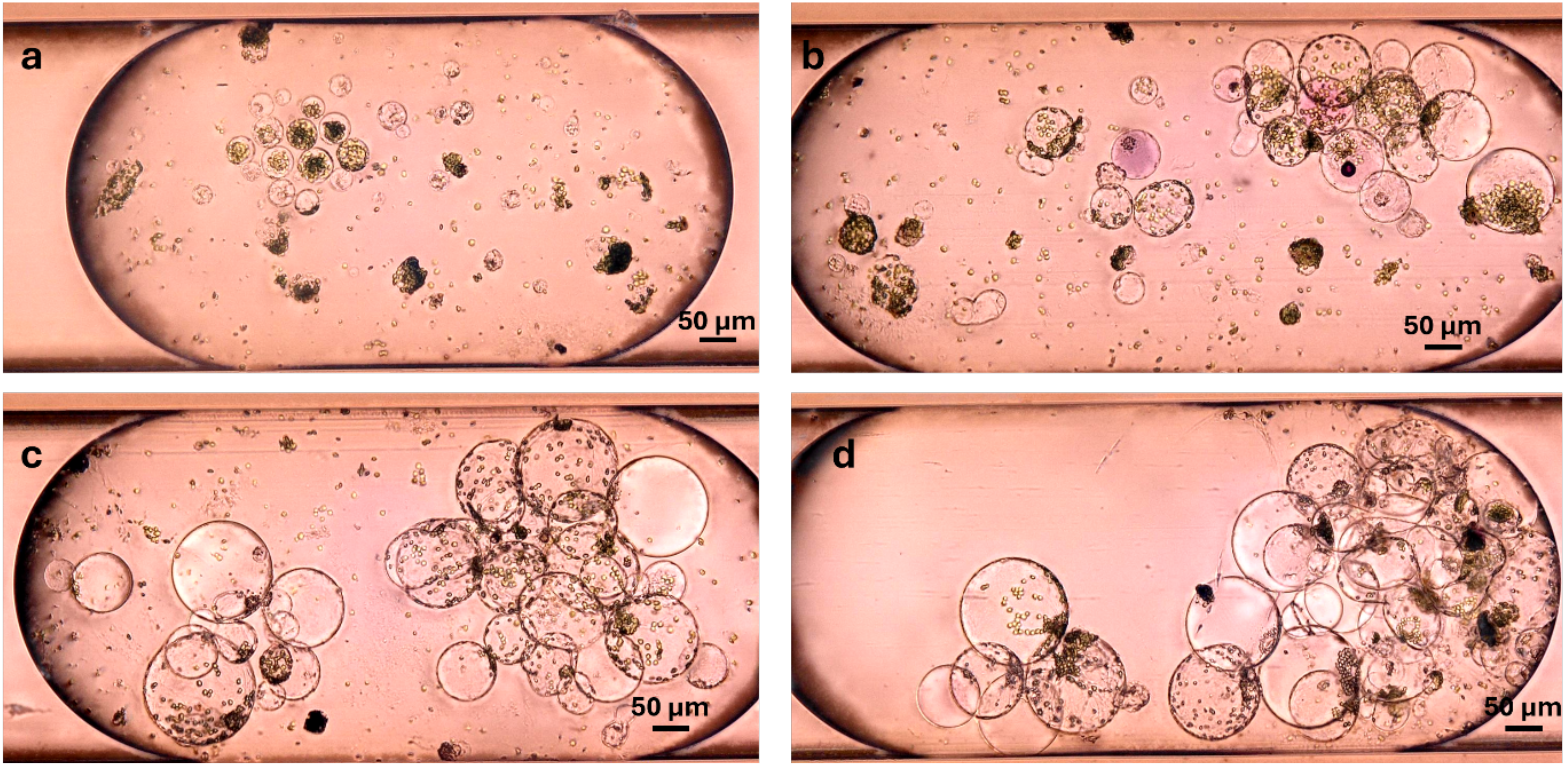
*Kalanchoe* leaf protoplasts at different time points during droplet cultivation: (a) day 0 – intact protoplasts introduced; some exhibit cytoplasmic condensation; (b) day 2 – cells increased in size and come into contact with each other, day 5 – further enlargement and first cell divisions observed; (d) day 7 – morphology remained largely unchanged from day 5. Images represent different droplets at each time point.

Importantly, despite the initial heterogeneity, *Kalanchoe* protoplasts within individual droplets were morphologically distinguishable and sufficiently separated to allow accurate identification, counting, and measurement of cell diameter over time. Based on these analyses, the size distribution of *Kalanchoe* protoplasts during cultivation was quantified (Fig. 6). Immediately after cell encapsulation, the average protoplast diameter in droplets was approximately 35 µm (range: 15–68 µm; *n* = 235), compared to 46 µm in parallel Petri dish cultures (*n* = 50). After two days, the droplet-cultured cells reached an average diameter of 49 µm (up to ~100 µm; *n* = 289), whereas Petri dish–cultured cells averaged 62 µm (*n* = 50). Peak growth in droplets was observed on day 5, with a maximum diameter of 167 µm and an average of 69 µm (*n* = 311), compared to 81 µm in Petri dishes (*n* = 50). Statistical analysis confirmed significant size differences across cultivation days (Kruskal–Wallis, χ^2^(3) = 191, p < .001), with post-hoc Dunn’s tests showing that all pairwise comparisons were significant (p < .001) except between day 5 and day 7 (p = 1.000). From day 5 to day 7, droplet-cultured protoplasts maintained a stable size distribution (average 68 µm; maximum 156 µm; *n* = 304), similar to the stabilization observed in Petri dishes (average 80 µm on day 7; *n* = 50).

**Figure 6.**
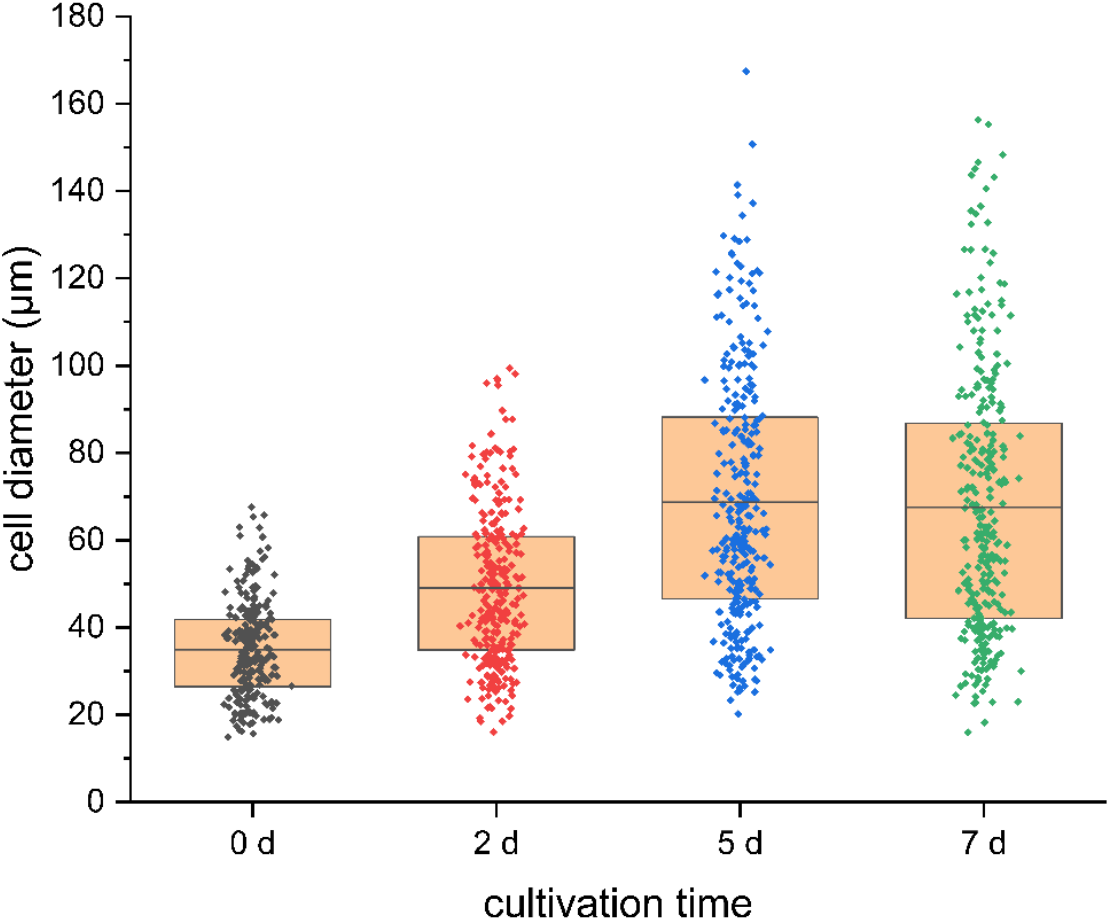
Size distribution of Kalanchoe protoplasts over time during droplet cultivation. Sample sizes (*n*): day 0 – 235, day 2 – 289, day 5 – 311, day 7 – 304; measured in 13 droplets for days 0–5 and 12 droplets for day 7. Significant size differences across days (Kruskal–Wallis, p < .001), with plateau between day 5 and 7.

Although only a limited number of cell divisions were observed, making it difficult to perform robust quantitative analysis of mitotic events, the continued increase in size of many protoplasts indicates ongoing metabolic activity and viability. Moreover, the observed variability in the fate of certain protoplast subpopulations is likely related to the differing regenerative potential of topophysical zones within the leaf [21], as in our study whole leaves were used as the source material for protoplast isolation. These findings suggest that selecting tissue from zones with higher regenerative capacity could improve the overall outcome of droplet-based cultivation.

### 3.3. Effect of low doses of growth regulators on protoplast survival during the first days of cultivation

Following successful cultivation under standard droplet conditions, tobacco protoplasts were selected for further exploration of the capabilities and advantages of the microfluidic platform. In this extended experiment, both the storage method of the tubes and the data acquisition strategy was modified, enabling tracking of specific droplets over time. This allowed observation of a much smaller and more defined group of protoplasts, in contrast to the broader population analyzed in previous experiments. As a test case, we examined the effects of plant growth regulators – 6-benzylaminopurine (BAP) and 1-naphthaleneacetic acid (NAA) – administered at significantly lower concentrations than the standard 1 mg·L^−1^, ranging from 0 to 150 µg·L^−1^. The aim was to determine the minimal concentration required to support protoplast viability during the initial days of cultivation. Figure 7 shows representative outcomes of protoplasts exposed to 0, 20, 50, 80, and 150 µg·L^−1^ NAA, with BAP applied at the same corresponding concentrations, as well as standard concentration of NAA and BAP for typical protoplasts culture of 1 mg·L^−1^. In the absence of growth regulators, protoplasts rapidly degenerated (Fig. 7a), whereas even low concentrations promoted survival and growth in the early stages (Fig. 7b–d). As reference, no degeneration and clear division of protoplasts by day 7 were observed in the protoplasts cultured in the presence of 1 mg·L^−1^ of NAA and BAP (Fig. 7e).

**Figure 7.**
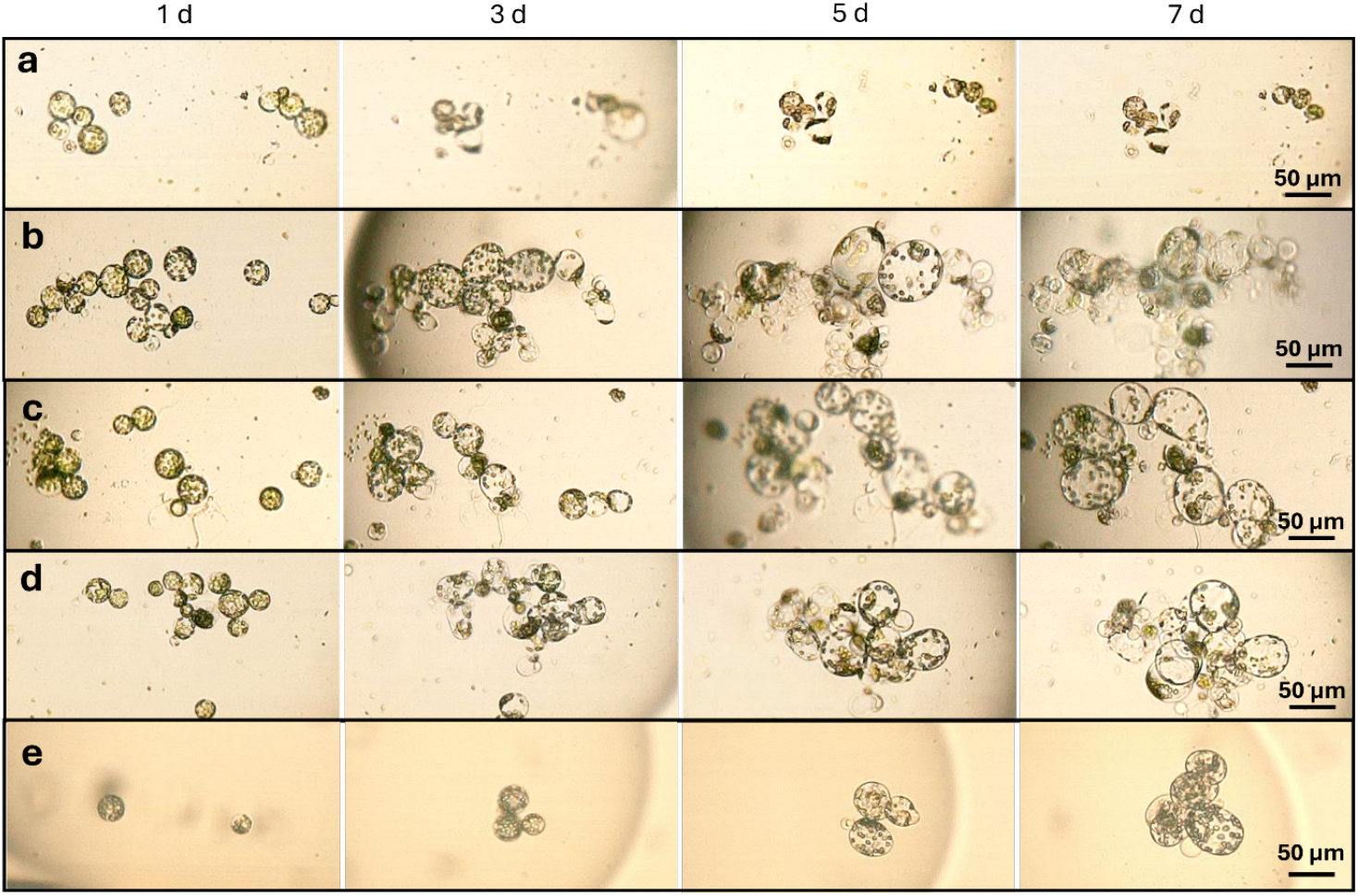
Development of tobacco protoplasts in droplets cultured with different concentrations of BAP and NAA: (**a**) 0 µg·L^−1^; (**b**) 50 µg·L^−1^; (**c**) 80 µg·L^−1^; (**d**) 150 µg·L^−1^, (**e**) 1 mg·L^−1^. Protoplasts treated with 1 mg·L^−1^ of growth regulators were photographed on days 0, 2, 4 and 7, respectively.

Although the applied growth regulator concentrations were generally insufficient to induce cell division, they did have a clear influence on cell viability. To quantify this effect, we monitored and counted morphologically healthy protoplasts in individual droplets throughout the cultivation period, normalizing these counts to the initial number of cells per droplet. Repeated measures ANOVA revealed significant effects of cultivation time (F(2.53, 244.9) = 33.4, p < 0.001) and growth regulator concentration (F(4, 97) = 475, p < 0.001) on the relative number of protoplasts, as well as a significant Time × Concentration interaction (F(10.1, 244.9) = 132.9, p < 0.001; Greenhouse–Geisser corrected). In the absence of growth regulators, protoplasts displayed a viability of approximately 35% by day 3, which dropped to 20% and 10% on days 5 and 7, respectively. In contrast, treatments with 20–150 µg·L^−1^ BAP + NAA promoted survival and proliferation, with the strongest responses observed from day 5 onwards at 80–150 µg·L^−1^, where cell numbers were maintained or increased relative to day 0 (Fig. 8a).

**Figure 8.**
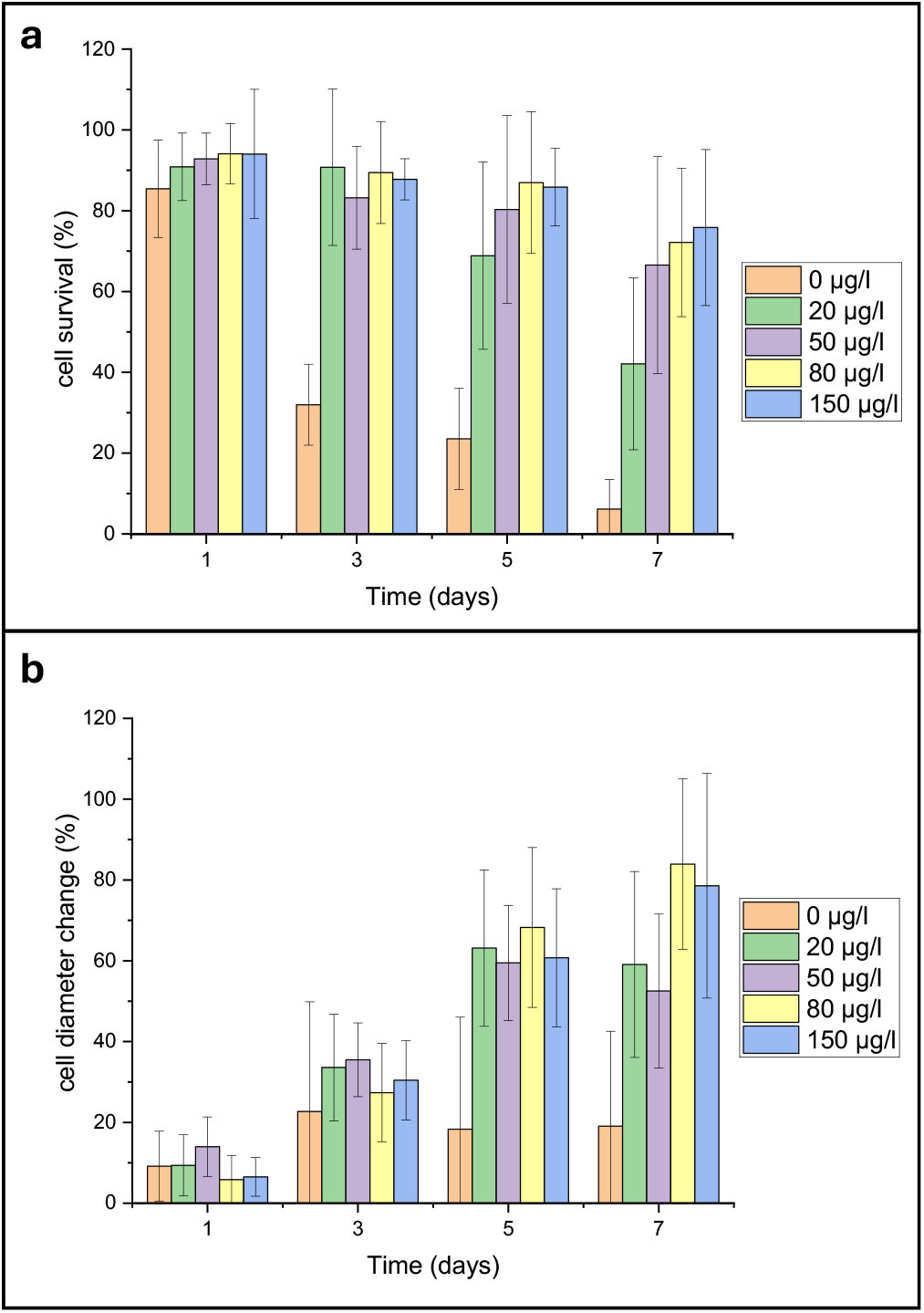
Characterization of tobacco protoplasts during droplet cultivation under varying concentrations of BAP and NAA (0, 20, 50, 80, and 150 µg·L^−1^): (**a**) Percentage of viable protoplasts over time, relative to day 0; (**b**) Change in average cell diameter during cultivation, normalized to the initial size at droplet loading. Initial sample sizes (*n*): 549 in 24 droplets (0 µg·L^−1^), 512 in 23 droplets (20 µg·L^−1^), 442 in 17 droplets (50 µg·L^−1^), 352 in 18 droplets (80 µg·L^−1^), and 414 in 18 droplets (150 µg·L^−1^). Error bars indicate standard deviation. Both protoplast number and size were significantly affected by time, concentration, and their interaction (repeated measures ANOVA, p < 0.001).

In addition to viability, we assessed changes in protoplast size over time. Repeated measures ANOVA also indicated significant effects of cultivation time (F(2.26, 191.9) = 237.2, p < 0.001) and concentration (F(4, 85) = 10.0, p < 0.001) on relative protoplast size, with a significant Time × Concentration interaction (F(9.03, 191.9) = 15.6, p < 0.001; Greenhouse–Geisser corrected). In the absence of phytohormones, surviving cells exhibited limited diameter increases – averaging only a 20% increase by day 3, with little to no further growth thereafter. In contrast, exposure to 20–150 µg·L^−1^ BAP + NAA induced progressive enlargement, with the most pronounced effect at 80–150 µg·L^−1^, where diameters nearly doubled relative to day 0 by day 7 (Fig. 8b). Estimated marginal means for each time × concentration condition are provided in *Supplementary Table S2* (cell number change) and *Supplementary Table S3* (cell size change). Although changes in cell diameter do not directly indicate viability or division potential, they reflect ongoing cellular activity and suggest that the protoplasts remained metabolically active under these droplet-based microfluidic conditions.

Although typical concentrations of growth regulators used in protoplast cultivation and plant regeneration studies are much higher, often ranging from 1 to 5 mg·L^−1^, we chose to investigate lower doses. Excessive concentrations of synthetic auxins and cytokinins are known to increase the risk of somaclonal variation, potentially leading to undesired genetic and phenotypic changes in regenerated plants [22]. By defining the minimal concentrations that still support protoplast survival and division, the risk of unwanted variation can be reduced. Additionally, using lower doses contributes to minimizing the environmental footprint of in vitro culture protocols [23,24].

## 4. Conclusion

In this study, we demonstrated the feasibility of using a droplet-based microfluidic platform for the cultivation of plant protoplasts. The platform allowed for cell encapsulation, maintenance, and observation of protoplasts across multiple species. Notably, species-specific differences in response were evident: tobacco leaf protoplasts showed the highest robustness and developmental potential, while mustard leaf protoplasts were highly sensitive to encapsulation-related stress and did not survive extended cultivation in droplets.

The microfluidic setup enables precise tracking of individual cells or defined subpopulations over time and allowed detailed investigation of early cellular responses to phytohormones. Supplementation with low concentration of cytokinin (BAP) and auxins (NAA) significantly enhanced early protoplast survival and growth, without additional benefit observed at higher concentration. These results highlight the sensitivity of the system for dose-response analysis.

Together, our findings demonstrate the potential of droplet-based microfluidics for protoplast-based assays and lay the foundation for future applications in plant developmental biology, regenerative research, and synthetic biology.

## Acknowledgements

Mrs. Marczakiewicz-Perera gratefully acknowledges financial support from Thuringian State Graduate Support. Dr. J. Cao gratefully acknowledges financial support from the Allianz für Industrie und Forschung (AiF) through the ZIM program, project “µProPlant” (FKZ: KK5240405AJ2). Helpful support and discussions with Dr. Welsch and Dr. Dovzhenko are gratefully acknowledged.

## Author contributions

PM: experiments conduction. PM, TW, MK, JC: Conceptualization and writing original draft.

## Data availability statement

The datasets generated during and/or analyzed during the current study are available from the corresponding author on reasonable request.

