## Supplemental Table for "Droplet-based microfluidics platform for investigation of protoplast development of three exemplary plant species"

Supplementary information for the research paper 'Droplet-based microfluidics platform for investigation of protoplast development of three exemplary plant species'

Paulina Marczakiewicz-Perera [a], Traud Winkelmann [b], Michael Köhler [a], Jialan Cao* [a]

*Supplementary Table S1.* Media composition and solutions sources used during protoplasts isolation and culture.

| Solutions | Manufacturer | Media | | | | | | |
|---|---|---|---|---|---|---|---|---|
| | | Preplasmolysis solution | BNE9 | Washing solution | Culture medium 8pm7 | MMM | F-PIN | Culture medium F-PCN |
| Sorbitol | Duchefa | 0.3 M | | | 0.25 $g{\cdot}L^{-1}$ | | | |
| Glycine | Duchefa | 0.1 M | | | | | | |
| $CaCl_2{\cdot}2H_2O$ | VWR | 0.05 M | 0.6 $g{\cdot}L^{-1}$ | 44.2 $g{\cdot}L^{-1}$ | | | 0.64 $g{\cdot}L^{-1}$ | |
| $NH_4NO_3$ | Grüssing | | 0.6 $g{\cdot}L^{-1}$ | 0.16 $g{\cdot}L^{-1}$ | 1.65 $g{\cdot}L^{-1}$ | | | |
| $KH_2PO_4$ | VWR | | 0.17 $g{\cdot}L^{-1}$ | 0.136 $g{\cdot}L^{-1}$ | 0.17 $g{\cdot}L^{-1}$ | | 0.17 $g{\cdot}L^{-1}$ | |
| KCl | Roth | | 0.3 $g{\cdot}L^{-1}$ | 18.8 $g{\cdot}L^{-1}$ | 0.3 $g{\cdot}L^{-1}$ | | | |
| Sucrose | Duchefa | | Approximately 125 $g{\cdot}L^{-1}$ | 23 $g{\cdot}L^{-1}$ | Approximately 125 $g{\cdot}L^{-1}$ | | Approximately 130 $g{\cdot}L^{-1}$ | Approximately 20 $g{\cdot}L^{-1}$ |
| Glucose | Applichem | | | | | | | 80 $g{\cdot}L^{-1}$ |
| $KNO_3$ | Merck | | 1.9 $g{\cdot}L^{-1}$ | | 1.9 $g{\cdot}L^{-1}$ | | 1.012 $g{\cdot}L^{-1}$ | 1.012 $g{\cdot}L^{-1}$ |
| $MgSO_4{\cdot}7H_2O$ | VWR | | 0.6 $g{\cdot}L^{-1}$ | | 3.7 $g{\cdot}L^{-1}$ | 1.25 $g{\cdot}L^{-1}$ | 0.37 $g{\cdot}L^{-1}$ | 0.37 $g{\cdot}L^{-1}$ |
| $MgCl_2{\cdot}6H_2O$ | VWR | | | | | 1.02 $g{\cdot}L^{-1}$ | | |
| KI | Roth | | | | 0.83 $mg{\cdot}L^{-1}$ | | 0.83 $mg{\cdot}L^{-1}$ | 0.83 $mg{\cdot}L^{-1}$ |
| $H_3BO_3$ | Roth | | | | 6.2 $mg{\cdot}L^{-1}$ | | 6.2 $mg{\cdot}L^{-1}$ | 6.2 $mg{\cdot}L^{-1}$ |
| $MnSO_4{\cdot}4H_2O$ | Geyer | | | | 26.6 $mg{\cdot}L^{-1}$ | | 26.6 $mg{\cdot}L^{-1}$ | 26.6 $mg{\cdot}L^{-1}$ |
| $ZnSO_4{\cdot}7H_2O$ | VWR | | | | 8.6 $mg{\cdot}L^{-1}$ | | 8.6 $mg{\cdot}L^{-1}$ | 8.6 $mg{\cdot}L^{-1}$ |
| $Na_2MoO_4{\cdot}2H_2O$ | ThGeyer Chemsolute | | | | 0.25 $mg{\cdot}L^{-1}$ | | 0.25 $mg{\cdot}L^{-1}$ | 0.25 $mg{\cdot}L^{-1}$ |
| $CuSO_4{\cdot}5H_2O$ | Merck | | | | 25 $\mu g{\cdot}L^{-1}$ | | 25 $\mu g{\cdot}L^{-1}$ | 25 $\mu g{\cdot}L^{-1}$ |
| $CoCl_2$ | Acros | | | | 13.5 $\mu g{\cdot}L^{-1}$ | | 13.5 $\mu g{\cdot}L^{-1}$ | 13.5 $\mu g{\cdot}L^{-1}$ |

| | | | | | | | | |
|---|---|---|---|---|---|---|---|---|
| NaFeEDTA | Roth | | | | 36.7 mg·L$^{-1}$ | | 36.7 mg·L$^{-1}$ | 36.7 mg·L$^{-1}$ |
| Myo-inositol | Merck | | | | 100 mg·L$^{-1}$ | | 200 mg·L$^{-1}$ | 200 mg·L$^{-1}$ |
| Nicotinic acid | Duchefa | | | | 2.5 mg·L$^{-1}$ | | 2 mg·L$^{-1}$ | 2 mg·L$^{-1}$ |
| Thiamine-HCl | Duchefa | | | | 10 mg·L$^{-1}$ | | 1 mg·L$^{-1}$ | 1 mg·L$^{-1}$ |
| Pyridoxine-HCl | Duchefa | | | | 1 mg·L$^{-1}$ | | 2 mg·L$^{-1}$ | 2 mg·L$^{-1}$ |
| Na-Pyruvate | Biochrom | | | | 0.02 g·L$^{-1}$ | | | |
| Biotin | Amresco | | | | | | 20 µg·L$^{-1}$ | 20 µg·L$^{-1}$ |
| Ca-pantothenate | Applichem | | | | | | 2 mg·L$^{-1}$ | 2 mg·L$^{-1}$ |
| Citric acid | VWR | | | | 0.04 g·L$^{-1}$ | | | |
| DL-Malic-acid | Duchefa | | | | 0.04 g·L$^{-1}$ | | | |
| Fumaric acid | Sigma Aldrich | | | | 0.04 g·L$^{-1}$ | | | |
| $NH_4Cl$ | Merck | | | | | | 1.06 g·L$^{-1}$ | 1.06 g·L$^{-1}$ |
| KOH | Merck | | | | | | 2.24 g·L$^{-1}$ | 2.24 g·L$^{-1}$ |
| Succinic acid | Roth | | | | | | 2.36 g·L$^{-1}$ | 2.36 g·L$^{-1}$ |
| MES | Applichem | | | | | 0.195 g·L$^{-1}$ | 0.195 g·L$^{-1}$ | |
| Mannitol | Duchefa | | | | 0.25 g·L$^{-1}$ | 85 g·L$^{-1}$ | | |
| Fructose | Acros | | | | 0.25 g·L$^{-1}$ | | | |
| Ribose | Duchefa | | | | 0.25 g·L$^{-1}$ | | | |
| Xylose | Alfa Aesar | | | | 0.25 g·L$^{-1}$ | | | |
| Mannose | Duchefa | | | | 0.25 g·L$^{-1}$ | | | |
| Rhamnose | Millipore | | | | 0.25 g·L$^{-1}$ | | | |
| Cellobiose | Fluka | | | | 0.25 g·L$^{-1}$ | | | |

*Supplementary Table S2.* Estimated marginal means (EMMs) of relative protoplast number change across cultivation days (1, 3, 5, 7) and growth regulator concentrations (0, 20, 50, 80, and 150 µg·$L^{-1}$ BAP + NAA), with standard errors and 95% confidence intervals.

| | | | | 95% Confidence Interval | |
|---|---|---|---|---|---|
| **Concentration** | **Time** | **Mean** | **SE** | **Lower** | **Upper** |
| **0** | **Day 1** | -0.1463 | 0.0171 | -0.1802 | -0.1123 |
| | **Day 3** | -0.6808 | 0.0264 | -0.7333 | -0.6284 |
| | **Day 5** | -0.7658 | 0.0364 | -0.8381 | -0.6935 |
| | **Day 7** | -0.9383 | 0.0410 | -1.0196 | -0.8570 |
| **20** | **Day 1** | -0.0913 | 0.0175 | -0.1259 | -0.0567 |
| | **Day 3** | -0.0909 | 0.0270 | -0.1444 | -0.0373 |
| | **Day 5** | -0.3113 | 0.0372 | -0.3852 | -0.2374 |
| | **Day 7** | -0.5791 | 0.0419 | -0.6622 | -0.4961 |
| **50** | **Day 1** | 0.1400 | 0.0203 | 0.0997 | 0.1803 |
| | **Day 3** | 0.3541 | 0.0314 | 0.2918 | 0.4164 |
| | **Day 5** | 0.5935 | 0.0433 | 0.5076 | 0.6795 |
| | **Day 7** | 0.5253 | 0.0487 | 0.4287 | 0.6219 |
| **80** | **Day 1** | 0.0572 | 0.0197 | 0.0181 | 0.0964 |
| | **Day 3** | 0.2739 | 0.0305 | 0.2134 | 0.3344 |
| | **Day 5** | 0.6833 | 0.0421 | 0.5998 | 0.7668 |
| | **Day 7** | 0.8389 | 0.0473 | 0.7450 | 0.9328 |
| **150** | **Day 1** | 0.0645 | 0.0187 | 0.0274 | 0.1016 |
| | **Day 3** | 0.3040 | 0.0289 | 0.2466 | 0.3614 |
| | **Day 5** | 0.6070 | 0.0399 | 0.5278 | 0.6862 |
| | **Day 7** | 0.7865 | 0.0449 | 0.6974 | 0.8756 |

*Supplementary Table S3.* Estimated marginal means (EMMs) of relative protoplast diameter change across cultivation days (1, 3, 5, 7) and growth regulator concentrations (0, 20, 50, 80, and 150 μg·L$^{-1}$ BAP + NAA), with standard errors and 95% confidence intervals.

| | | | | 95% Confidence Interval | |
|---|---|---|---|---|---|
| **concentration** | **Time** | **Mean** | **SE** | **Lower** | **Upper** |
| **0** | **Day 1** | 0.1167 | 0.0203 | 0.0763 | 0.1571 |
| | **Day 3** | 0.2717 | 0.0417 | 0.1888 | 0.3545 |
| | **Day 5** | 0.2008 | 0.0565 | 0.0885 | 0.3132 |
| | **Day 7** | 0.1892 | 0.0669 | 0.0561 | 0.3222 |
| **20** | **Day 1** | 0.0930 | 0.0147 | 0.0639 | 0.1222 |
| | **Day 3** | 0.3357 | 0.0301 | 0.2758 | 0.3955 |
| | **Day 5** | 0.6317 | 0.0408 | 0.5506 | 0.7129 |
| | **Day 7** | 0.5909 | 0.0483 | 0.4948 | 0.6870 |
| **50** | **Day 1** | 0.1400 | 0.0171 | 0.1061 | 0.1739 |
| | **Day 3** | 0.3541 | 0.0350 | 0.2845 | 0.4237 |
| | **Day 5** | 0.5935 | 0.0475 | 0.4991 | 0.6879 |
| | **Day 7** | 0.5253 | 0.0562 | 0.4135 | 0.6371 |
| **80** | **Day 1** | 0.0572 | 0.0166 | 0.0242 | 0.0902 |
| | **Day 3** | 0.2739 | 0.0340 | 0.2063 | 0.3415 |
| | **Day 5** | 0.6833 | 0.0461 | 0.5916 | 0.7751 |
| | **Day 7** | 0.8389 | 0.0546 | 0.7302 | 0.9475 |
| **150** | **Day 1** | 0.0645 | 0.0157 | 0.0332 | 0.0958 |
| | **Day 3** | 0.3040 | 0.0323 | 0.2398 | 0.3682 |
| | **Day 5** | 0.6070 | 0.0438 | 0.5200 | 0.6940 |
| | **Day 7** | 0.7865 | 0.0518 | 0.6834 | 0.8896 |
